# Efficient assembly and long-term stability of defensive microbiomes via private resources and community bistability

**DOI:** 10.1101/457721

**Authors:** Gergely Boza, Sarah F. Worsley, Douglas W. Yu, Istvan Scheuring

**Author notes:** /1706; Address: Klebelsberg Kunó str. 3, H-8237 Tihany, Hungary.

## Abstract

Understanding the mechanisms promoting the assembly and maintenance of host-beneficial microbiomes is an open problem. An increasing amount of evidence supports the idea that animal and plant hosts can use ‘private resources’ and the ecological phenomenon known as ‘community bistability’ to favour some microbial strains over others. We briefly review empirical evidence showing that hosts can: (i) protect the growth of beneficial strains in an isolated habitat, (ii) use antibiotic compounds to suppress non-beneficial, competitor strains, and (iii) provide resources (for a limited time) that only beneficial strains are able to translate into an increased rate of growth, reproduction, or antibiotic production. We then demonstrate in a spatially explicit, individual-based model that these three mechanisms act similarly by selectively promoting the initial proliferation of preferred strains, that is, by acting as a private resource. By explicitly modelling localized microbial interactions and diffusion dynamics, we further show that an intermediate level of antibiotic diffusion is the most efficient mechanism in promoting preferred strains and that that there is a wide range of conditions under which hosts can promote the assembly of a self-sustaining defensive microbiome. This, in turn, supports the idea that hosts readily evolve to promote host-beneficial defensive microbiomes.

## Introduction

A growing number of studies show that microbiome composition is structured by competition [1, 2, 3, 4, 5, 6, 7], and it is hypothesized that a host could evolve to bias these processes to promote the establishment of host-beneficial microbes [6, 8, 9, 10, 11, 12, 13]. Indeed, such microbes need support because, first, it is inherently difficult to establish a colony of host-beneficial microbes in the face of competition against the huge pool of available host-neutral or host-harmful species [1, 14, 15, 16, 17], and second, these microbes often produce costly compounds that, although equipping them to be beneficial for the host, render them competitively inferior to non-beneficial, or even parasitic, microbes [18].

We distinguish three mechanisms by which a host can selectively favour beneficial strain(s), namely (1) *providing a habitable space* that the desired bacterial partner has preferential access to, (2) *production of specific compounds* by the host that selectively poison undesired bacteria, and (3) *providing a food resource* that the desired partner is better able to metabolise. We now briefly review examples of each:

1. *Providing a habitable space that the desired bacterial partner has preferential access to.* — Vertical and pseudo-vertical transmissions fall into this category [1, 19, 20, 21, 22, 23]. In strict vertical transmission, host germline cells are infected with symbionts [22, 24]. Less strict transmission (‘pseudo-vertical’) is achieved by keeping non-colonised host offspring in isolation after birth until the parental microbiome can colonise it, which then shapes the composition of subsequent colonists from the environment [9, 11, 22]. In either case, the host ensures a ‘competitor-free space’ for inherited microbes, which are allowed time and resources to grow on a new-born host before being exposed to competition with other colonists. For example, newly emerging *Acromyrmex* leafcutter ants are inoculated with antibiotic-producing *Pseudonocardia* bacteria within a 24-hour window after hatching [8, 25]. Mature worker ants serve as the source, carrying *Pseudonocardia* on their propleural plates, which grow to high density around specialised exocrine glands that likely provide nutrients for bacterial growth [26, 27] (thus also serving as an example of a resource that can be metabolized by the preferred bacteria, discussed in 3. below). Similarly, female beewolf digger wasps (*Philanthus*, *Philanthinus*, *Trachypus*) inoculate their brood cell walls with species of *Streptomyces* that they maintain in their antennal glands [28, 29, 30]. These bacteria become directly incorporated into the larval cocoon, where they dominate and produce an array of antibiotics that protect the developing larva against infection [29, 30, 31]. Analogous to the above examples, the agricultural process of applying bacteria, such as antibiotic-producing *Pseudomonas* and nitrogen-fixing *Rhizobia*, to crop seeds before sowing mimics pseudo”vertical”transmission, by ensuring that high densities of beneficial bacteria have better access to root exudates and are favoured during establishment on the plant [32, 33]. Priority effects have also been demonstrated for mycorrhizae [34], bees [35, 36, 37, 38], wasps [28], leafcutter ants [25, 39], birds [40], plants [41], and humans [42].
2. *Hosts producing compounds poisoning non-desired bacteria, whilst allowing desired strains to grow.* — A wide range of plants secrete compounds, known as allelochemicals, that have toxic effects on a broad range of bacteria, fungi, and invertebrates in the rhizosphere, as well as other plants growing nearby [43, 44, 45, 46]. For example, the compound DIMBOA is an antimicrobial produced by maize seedlings [45], which the plant-beneficial species *Pseudomonas putida* is able to degrade, thus avoiding its effects. *P. putida* also uses this compound as a chemoattractant and a signal for upregulating the production of the broad-spectrum antibiotic phenazine [45]. Together, these mechanisms allow *P. putida* to colonise maize roots in the presence of mostly DIMBOA-intolerant, competitor bacteria [45]. Similarly, the rhizobial species, *Mesorhizobium tianshanense*, which forms root nodules on liquorice plants is able to outcompete other bacteria in the rhizosphere due to an efflux mechanism that confers resistance to the antimicrobial compound canavanine. Canavanine is abundant in liquorice root exudates and thus allows the host to filter out non-beneficial rhizobial species [47]. As another example, nitric oxide (NO), a potent oxidising agent and antimicrobial, can play an important role in dictating symbiont specificity [48, 49]. A classic example arises during the symbiosis between the bobtail squid, *Euprymna scolopes*, and bioluminescent bacteria in the species *Vibrio fischeri. V. fischeri* are the exclusive colonisers of the squid’s light organ, where they emit light to deceive predators, and are acquired horizontally from the environment within 48 hours after squid hatching [50]. High nitric-oxide synthase (NOS) activity and its product NO can be detected in the epithelial mucus of the light organ during the early stages of bacterial colonisation [51], which *V. fischeri* are able to tolerate via the activity of two proteins, flavohemoglobin (*Hmp*) and a heme NO/oxygen-binding protein (H-NOX) [52, 53, 54, 55]. Eliminating the genes for these proteins in *V. fischeri* leads to colonisation deficiency [53, 55], and diminishing the concentration of host NO results in a greater diversity of non-mutualistic bacterial species in the light organ epithelium [51]. Similar mechanisms of host selection are reported for animals as well. For example, members of the *Hydra* family produce antibacterial arminins that help them to shape the establishment of the bacterial microbiota during their embryogenesis [56]. *Hydra* not only suppress the undesired strains [56] but also modify the quorum-sensing signals through which bacteria communicate, hence manipulating the social behaviour of bacteria [57].
3. *A food resource that the desired partner is better able to metabolise.* — Enhanced metabolic activity from consuming a private resource can confer competitive superiority to a preferred microbial strain. Besides acquiring higher reproduction and growth rates, the beneficial bacteria can also achieve a higher rate of antibiotic production, resulting in the suppression of competitors [58], or a higher production of other factors that promote colonization and symbiotic interaction with the host, such as adhesive molecules facilitating biofilm formation on the host surface [59, 60]. The provision of specific metabolites is thought to play a key role in structuring the species-specific microbial communities associated with marine corals [61, 62]. Coral juveniles, as well as their dinoflagellate symbionts, produce large quantities of the compound dimethylsulfoniopropionate (DMSP) [63]. *In vitro* and metagenomic studies have shown that several coral-associated bacterial groups can specifically metabolise the DMSP and use it as a sole carbon and sulphur source [61, 62, 64]. Such species are also amongst the first bacteria to colonise coral larvae, suggesting a nutritional advantage for them over bacteria that cannot degrade DMSP [61, 65]. This includes a species of *Pseudovibrio* which can additionally use DMSP as a precursor for the production of antibiotics which inhibit coral pathogens [62]. Another example of a specific host-derived resource is human breast milk, which is known to contain a large number of complex oligosaccharides that are preferentially consumed by a single species of co-adapted gut bacterium *Bifidobacterium longum* subsp. *infantis* [66].

In plants, experiments have shown that root exudates can be directly metabolised by the microorganisms that live endophytically within the plant roots [67, 68, 69]. Different species exude different groups of metabolites, and studies suggest that plant hosts may be able to tailor root exudate composition in order to recruit bacteria with particular metabolic traits [43, 67, 70]. For example, the concentration of the plant phytohormone salicylic acid (SA) has been shown to correlate with the abundance of several bacterial taxa, including the antibiotic-producing genus *Streptomyces* [70, 71], which can use SA as a sole carbon source [71, 72]. As discussed earlier, leaf-cutter ants exocrine glands which provide a nutrient source for *Pseudonocardia* bacterial growth also fall into this category [26].

These mechanisms achieve one of two effects: (i) they either ensure the protected growth of the preferred strains and/or (ii) they enhance the competitive abilities of preferred strains against non-preferred strains for certain duration of time. Essentially, a *host can use multiple means to provide a ‘private resource’, in the form of space and/or food, to a subset of bacterial strains*, and if those strains are beneficial to the hosts, the host is selected to evolve and apply one or more of these mechanisms to assemble host-beneficial microbiomes.

We now abstract these mechanisms into an individual-based, spatially-explicit model of host-associated defensive microbiomes (reviews in 9, 29, 30), which typically contain antibiotic-producing, and at the same time resistant, bacteria [73, 74]. In our model, dispersal and *direct competition for empty sites* is limited to small numbers of neighbouring individuals, in accordance with experimental results [75]. At the same time, due to diffusion, we allow *indirect, antibiotic-mediated competition* to occur amongst distant bacteria. We aim to understand the conditions under which host species can promote the growth and dominance of antibiotic-producing, defensive microbiomes.

## Materials and methods

We are interested in how the host influences the population dynamics of two different bacterial strains: an antibiotic-producing, resistant beneficial strain (**B**), and a non-producing, sensitive parasitic strain (**P**) (Fig. 1a). We model the host implicitly by assuming it is able to manipulate the composition of its microbiome through resource supply on its surface, upon which colonising individual bacteria compete for space with their neighbours according to their reproduction rates. The host surface further serves as a medium for spatially limited diffusion of the antibiotic. For this, we employ an individual-based model in which we model the host surface as a rectangular grid with toroidal boundary conditions (*N* = *M* * *M*) serving as the habitat for coloniser bacteria (Fig. 1b). Each grid point can be empty or inhabited by a single individual, and interactions take place within the immediate neighbourhood of the focal grid point. Time is measured in units of update steps. We assume that the dynamics of cell reproduction and death processes are much slower than the small-molecule dynamics, so the cell populations are updated after *u* (*u* ≫ 1) update steps in antibiotic dynamics, during which the whole grid is updated in all relevant intracellular and extracellular processes related to the small-molecule (antibiotic) dynamics (*N* number of sites). In the cellular update steps, *εN* number of randomly chosen grid cells are updated in the birth and death processes, and *ε* is selected to be a small positive number (see Supplementary information 1).

**Figure 1.**
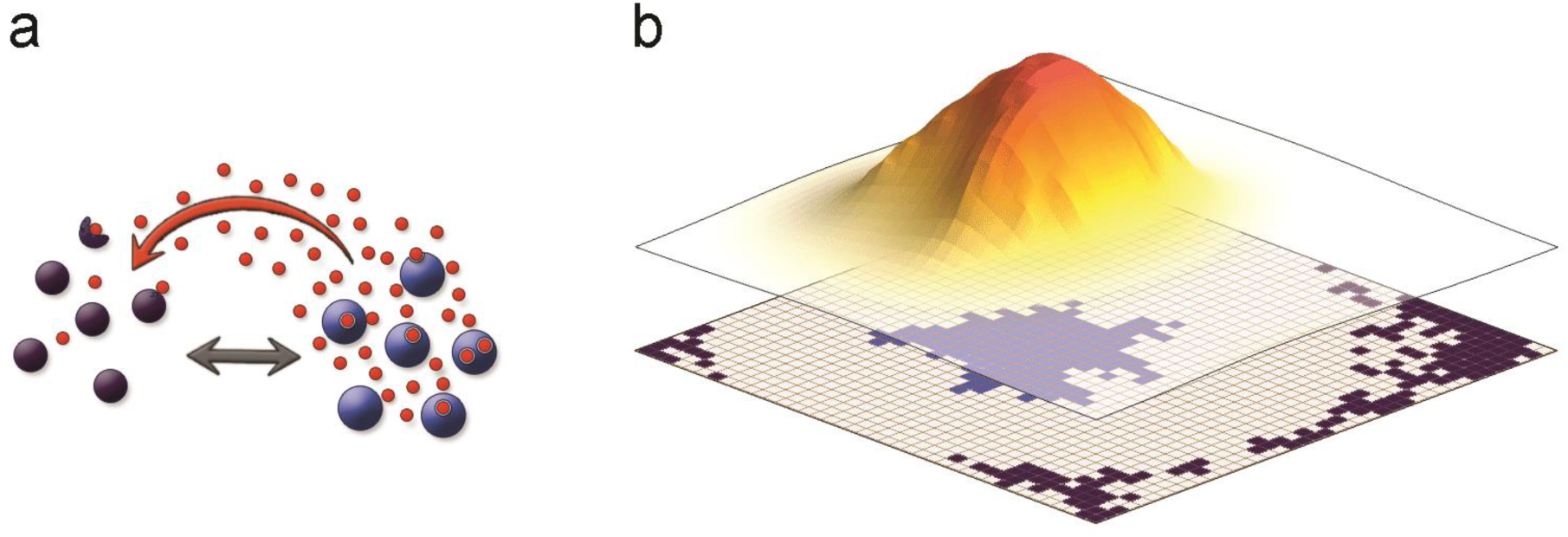
**a** We model two types of strains, parasitic (violet shading) and antibiotic-producer (blue shading), which compete with each other directly (grey arrow), and indirectly via the diffusing antibiotic (red dots and coloured arrows). **b** The modelled *N* = *M* * *M* grid (bottom layer) represents the colonisable surface of the host, and each point in the grid can be inhabited by a single individual (coloured quadrant). The produced antibiotic (upper layer) diffuses freely on the grid, and its concentration decreases farther from the producing source (the shading and height depicting the concentration) and also decays with time. Relevant model parameters are: *D* = 5, *Δt* = 1/10, *u* = 100, for **a** *n*_B,0_ = 100, *N* = 10 000, and for **b** *n*_B,0_ = 1, *M* = 40, *ρ* = 1, *α*_B_ = 0.5, *β*_B_ = 0.6, *γ*_B_ = 0.3, *φ* = 0.5.

The private resource(s) provided by the host can confer two kinds of benefits to the beneficial strain. We call the first kind **Protected Growth** (recall mechanisms 1 and 2 from *Introduction*), because *the parasitic strain is prevented from invading (i.e. colonising) the host until time τ*. Accordingly, **B** is given preferred access to host-provided food or space or is solely resistant to host-produced allelochemicals until time *τ*, after which time the host resource is made ‘public’ by giving the parasitic strain access to host-provided food or space or by withdrawing the host-produced compounds facilitating **B** or poisoning **P**. We call the second kind **Enhanced Metabolism**, because *although* **P** *is allowed to invade starting from time* 0, ***B’****s metabolism is enhanced until time τ*, *after which this enhancement lapses* (recall mechanism 3 from *Introduction*). The simplest outcome of enhanced metabolism is that **B**’s advantage in metabolising host-provided food causes its population growth rate to be increased by an amount of *r*_B,pr_(*t*) until time *τ*, after which *r*_B,pr_(*t*) = 0 (e.g. *r*_B,pr_(*t*) ≥ 0|*t* < *τ* and *r*_B,pr_(*t*) = 0|*t* ≥ *τ*), where index pr denotes the private resource. An alternative outcome is that **B** is able to use the host-provided food to *increase its own antibiotic production rate* (*ρ*_B_(*t*) = *ρ*_B,pr_(*t*) + *ρ*_B,0_), without incurring higher unit costs. Thus, similarly as above, we distinguish a higher production rate fuelled by host-provided resource (*ρ*_B,pr_(*t*) ≥ 0|*t* < *τ*), and a lower, baseline production rate when the resource is not supplied after time *τ* (*ρ*_B,pr_(*t*) = 0|*t* ≥ *τ*). Naturally, *ρ*_B,0_ > 0, while the production rate of the non-producing strain is always zero (*ρ*_P_(*t*) = 0). (Alternatively, but not modelled here, the resource could allow the antibiotic to be effective at a lower threshold concentration before *τ* and at higher level after *τ*, which would give similar results to the previous).

### Dynamics of the antibiotic molecules

The beneficial strain produces and exports antibiotic at rate *ρ*_B_. into the extracellular environment, resulting in a distribution of concentrations *A*^Ext^(*i, t*) at position *i* at time *t*.

The molecules are taken up by the cells at rates *α*_B_ and *α*_P_ (*α*_B_ ≤ *α*_P_) by the **B** and the **P** strains, respectively, resulting in an *A*^Int^(*i,t*) interior concentration within the cell at position *i* at time *t*. The cells decompose the intracellular antibiotics at rates *γ*_B_ and *γ*_P_ (*γ*_B_ ≥ *γ*_P_), and they can also perform active outbound transport, i.e. controlled efflux, to release intracellular antibiotics at rates *β*_B_ and *β*_P_ (*β*_B_ ≥ *β*_P_). The antibiotics decays at rate *φ* in the environment.

The model implements the three major antibiotic-resistance mechanisms: (a) reduced influx through the membrane (*α*_B_), (b) a higher rate of intracellular decomposition and neutralisation (*γ*_B_), and (c) increased efflux of the molecules (*β*_B_), and combinations of these mechanisms [73, 76, 77, 78, 79].

Based on the continuous reaction-diffusion model of the detailed dynamics (see Supplementary information 2 for details) [80], the time-and-space-discretised dynamics of antibiotic concentration at site *i* on the grid and at time *t* + Δ*t* in the extracellular environment can be written as

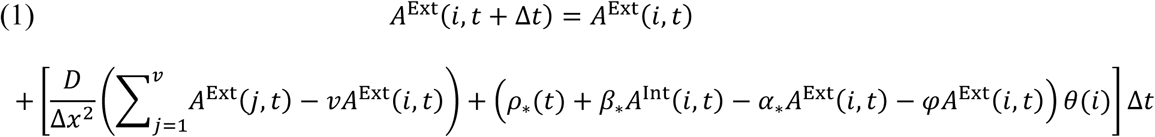

where the first term corresponds to the diffusion of antibiotics according to the discretised diffusion algorithm between the four nearest neighbouring points (*v* = 4) (Neumann-neighbourhood: north, south, east, west); Δ*x* is the spatial resolution, and Δ*t* is the time resolution. The diffusion rate of the antibiotics, *D*, is measured in the unit of *x*^2^/*t*, where *x* denotes the spatial resolution, here one cell of the grid, and *t* stands for time measured as an update step. *θ*(i) takes the value one if there is a cell at the site *i*, else being zero. The dynamics of intracellular concentration of the antibiotic at the site *i* can be written as

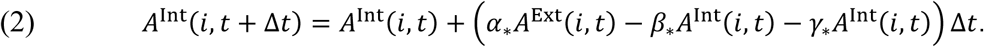

Naturally *A*^Int^(*i, t* + Δ*t*) = *A*^Int^(*i, t*) = *A*^Ext^(*i, t*) = 0 if there is no cell at site *i*.

### Growth dynamics of the cells

For the birth and death processes, we define the growth rate of the antibiotic-producing (**B**) and non-producing (**P**) strains respectively as

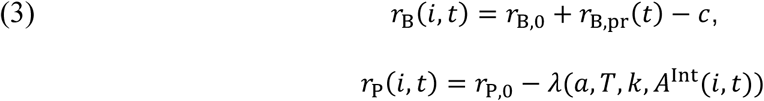

where *c* is the decrease in reproduction rate because of the costly processes of antibiotic production and resistance. The reproduction rates *r*_B,0_, *r*_P,0_, and *r*_B,pr_(*t*) correspond to normal (baseline) and temporarily increased resource conditions, respectively. The effect of the antibiotic *λ*(*α,T,k,A*^lnt^(*i,t*)) on the **P** strain’s reproduction rate depends on the critical threshold (*T*), the maximum effect (*a*), the steepness of the dosage effect (*k*), and the actual intracellular concentration of the antibiotic in the sensitive cell at the site *i* (*A*^Int^(*i, t*)). Following empirical observations [58], we define a general sigmoid function for the effect of the antibiotic:

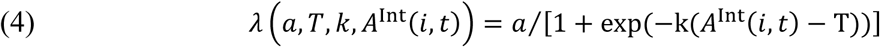

### Dynamics of the population

Population dynamics are represented by a death-birth process in which a randomly chosen focal individual at site *i* dies, and individuals from its Moore neighbourhood (8 nearest neighbours, *w* = 8) can reproduce and place a progeny into this focal empty site, with probability proportional to their growth rates

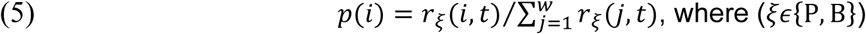

At the beginning of the simulation, the beneficial strain is represented in low numbers (*n*_B,0_), and the parasitic strain is missing (*n*_P,0_ = 0).

### Invasion tests

We carried out two sets of invasion tests to demonstrate how host-provided private resources can result in self-sustaining, beneficial microbiomes, even if the advantage provided by the resource eventually diminishes. In the first test, we used time, while in the second, we used colony size as the signal to switch from private to public resources, or in other words, to stop the host’s selective support for the beneficial strain.

#### Invasion test 1. Time-limited supply of private resources

We model the two kinds of benefits conveyed by the private resources, as discussed earlier, the (1) Protected Growth of the beneficial strain for *τ* time and (2) the Enhanced Metabolism of the beneficial strain for *τ* time, either leading to (2a) a higher population growth rate by the beneficial strain or to (2b) an increased antibiotic production by the beneficial strain.

In the Protected Growth scenario, before *τ*, an *s*_+_ proportion of cohesive space on the host surface (*s*_+_ = *ss*/*N* * 100, where *ss* is the number of protected sites) provides a safe growth opportunity for the beneficial strain, as individuals from the parasitic strain are prevented from invading (strict and pseudo-vertical transmission), or parasitic individuals invading this region get killed off (via host-provided allelochemicals). Only after *τ* time has passed is the parasitic strain allowed to gain a foothold anywhere on the grid (empty or occupied). In other words, the private space resource becomes public at time *τ*. During an invasion attempt, we place *n*_P,t_ = 0 number of individuals around a randomly selected focal grid point in a connected cluster with probability *f* in each time step (if there are empty places, subsequent individuals will be placed next to the focal site, but non-empty grid points can also be occupied if no empty places are available). In the Enhanced Metabolism scenario, the beneficial strain experiences increasing advantages of *r*_+_ = (*r*_B,pr_ (*t*) + *r*_B,0_ * 100 or *ρ*_+_ = (*ρ*_B,pr_(*t*) + *ρ*_B,0_)/*ρ*_B,0_ * 100 for *τ* time, respectively, and *n*_P,t_ = 0 number of parasitic-strain individuals are allowed to invade with probability *f* in each time step, starting from the beginning.

#### Invasion test 2. Protected Growth of the beneficial strain to a minimum colony size

Here, we let the host resource, the habitat, be private until the beneficial strain reaches a minimum colony size, which we call the Colony Size at Invasion (*CSI*). We only allow the parasitic strain to start invading empty places after the resident **B** strain’s colony size has grown to the *CSI* (*CSI* = *q*/*N* * 100, where *q* is the number of sites inhabited by **B**). The invasion proceeds with probability *f* and with *n*_P,t_ number of invaders until the grid is fully occupied by individuals. As a motivating example, one can think of a small host ‘crypt’ in which beneficial strain is initially be housed, but the strain eventually outgrows the crypt and colonises the host surface, at which point, the host can only provide resources in a way that makes them publicly available.

## Results

### Invasion test 1. Time-limited supply of private resources

As discussed in the *Introduction*, the host has multiple mechanisms by which it can provide private resources. We find that protecting initial growth (Fig. 2a, b), increasing the population growth rate (Fig. 2c, d), and/or enhancing the antibiotic effectiveness (Fig. 2e, f) of the beneficial strain can all result in a self-sustaining, beneficial-strain-dominated microbiome that is resistant to invasion even after the host resource is made public (at time *τ*) and the beneficial strain starts to experience a competitive disadvantage due to the costs of antibiotic production and of expressing its antibiotic-resistance traits. In all three scenarios, the longer the time *τ* that the resource is private (Fig. 2, x-axis), the less of an advantage, in the form of protected growth (here *τ* correlates with the size of the colony, see Supplementary information 1 Fig. 1), increased population growth (*r*_+_), or increased antibiotic production (*ρ*_+_) (y-axis), is required for the beneficial strain to be able to resist invasion after the resource becomes public. This is because invasion resistance is dependent on the beneficial colony reaching a sufficiently large size and on the concentration of antibiotic the colony produces and transports into the environment.

**Figure 2.**
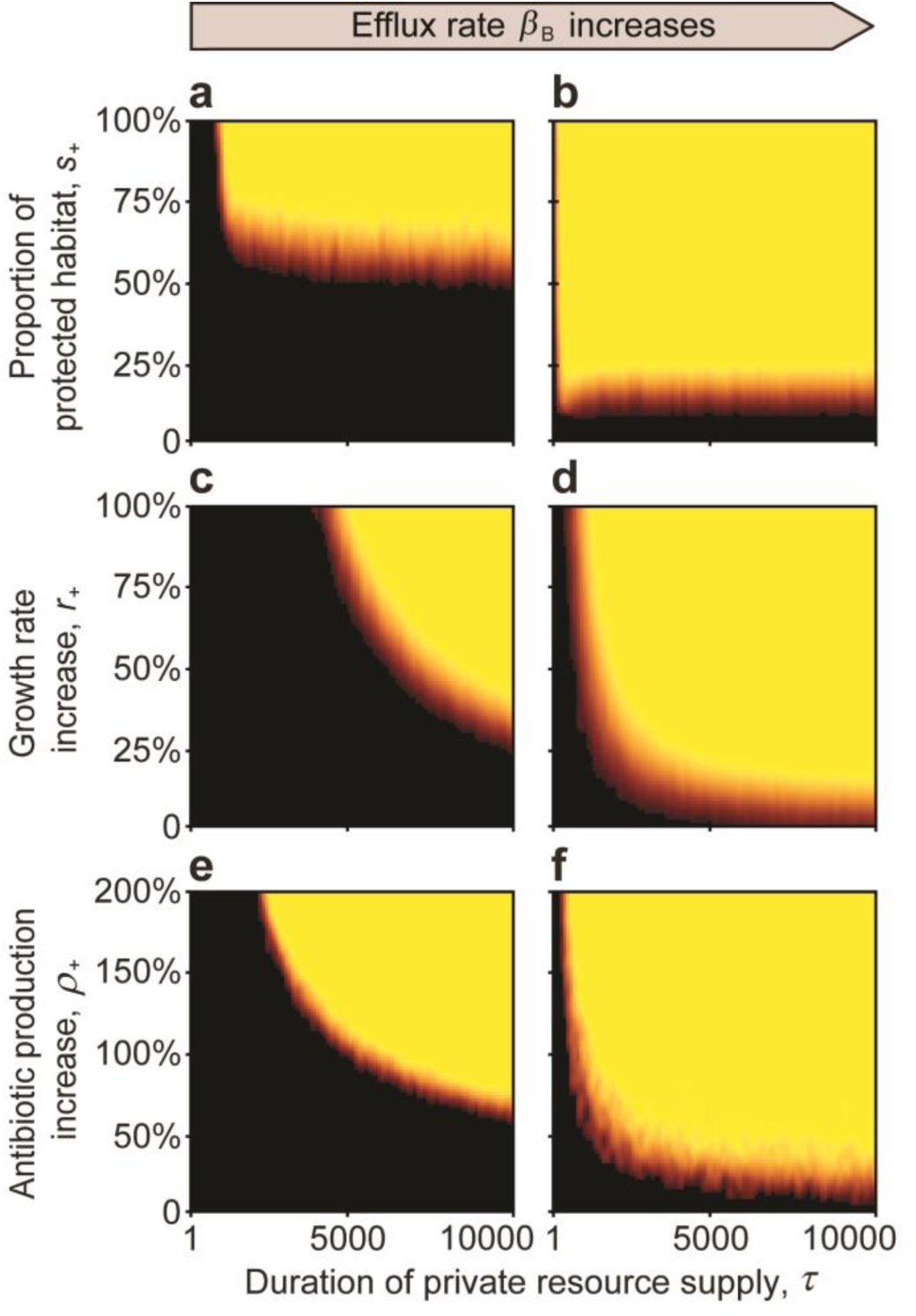
The effect of a private resource supplied by the host for a limited time *τ* (*Invasion Test 1*). Black areas indicate parameter space where the non-producing parasitic strain can invade, and yellow indicates that the beneficial strain is able to resist invasion. Orange to red colours indicate mixed outcomes. In general, the beneficial strain (yellow) dominates over a larger proportion of the parameter space as the duration of the private resource supply lengthens, regardless of whether the beneficial strain enjoys outright protected growth (**a, b**), an increased rate of population growth (**c, d**), or an increased rate of antibiotic production (e, f). The efflux of accumulated intracellular antibiotic in the antibiotic-producing beneficial strain also aids beneficial-strain dominance (*β*_B_ = 0 for **a, c, e,** and *β*_B_ = 0.25 for **b, d, f**). Simulations were run with 5 replicates for 100 000 generations or until the population was homogenous. Model parameters are: *r*_B,0_ = 0.8, *r*_P,0_ = 0.8, *c* = 0.1, *ρ*_B,0_ = 1, *α*_B_ = 0.5, *α*_P_ = 0.5, *β*_P_ = 0, *γ*_B_ = 0.4, *γ*_P_ = 0.4, *D* = 5, *a* = 1, *T* = 1, *k* = 25, *N* = 10 000, *n*_B,0_ = 100, *n*_P,t_ = 10, *f* = 0.01, *D* = 5, Δ*t* = 1/10, *u* = 100, *φ* = 0.3 for **a, b, c, d,** *φ* = 0.4 for **e, f,** and *r*_B,pr_(*t*) = 0, and *ρ*_B,pr_(*t*) = 0 when applicable.

We also observe that if the physiological mechanism of resistance by the beneficial strain to its own antibiotic is efflux, this can additionally enhance invasion resistance, even if the supply time is short and the advantage conferred by the private resource is small (Fig. 2a, b, c vs. e, d, f). The reason is that re-exporting any ingested antibiotic increases the environmental concentration of antibiotic, which aids suppression of invading parasitic strains.

### Invasion test 2. Protected Growth of the beneficial strain until a minimum colony size

Consistent with the results from Invasion test 1, if the beneficial colony reaches a critical size, (the Minimum Sustainable Colony size: *MSC*) it becomes resistant to invasion over a wide range of parameters after the private resource is made public (Fig. 3). Again, having antibiotic efflux as the resistance mechanism promotes invasion resistance (Fig. 3 and 4), whereas (and intuitively) a higher rate of extracellular decay of antibiotic counteracts this effect (Fig. 3 and 4). When a large amount of antibiotic is in the environment, because efflux is high and decay is low (Fig. 3a and Fig. 4a, c), the beneficial strain is able to dominate over a wide range of diffusion rates. However, when efflux is weak and the extracellular decay rate is high, only high diffusion rates allow the beneficial strain to dominate (Fig. 4d). This is because at low diffusion rates, the antibiotic produced in the centre of the colony is lost due to decomposition before reaching the colony edge by diffusion, where it would have attacked invaders. In contrast, at high diffusion rates, more of the resident colony’s antibiotic production is recruited to fight invasion (Fig. 3, 4, and 5).

**Figure 3.**
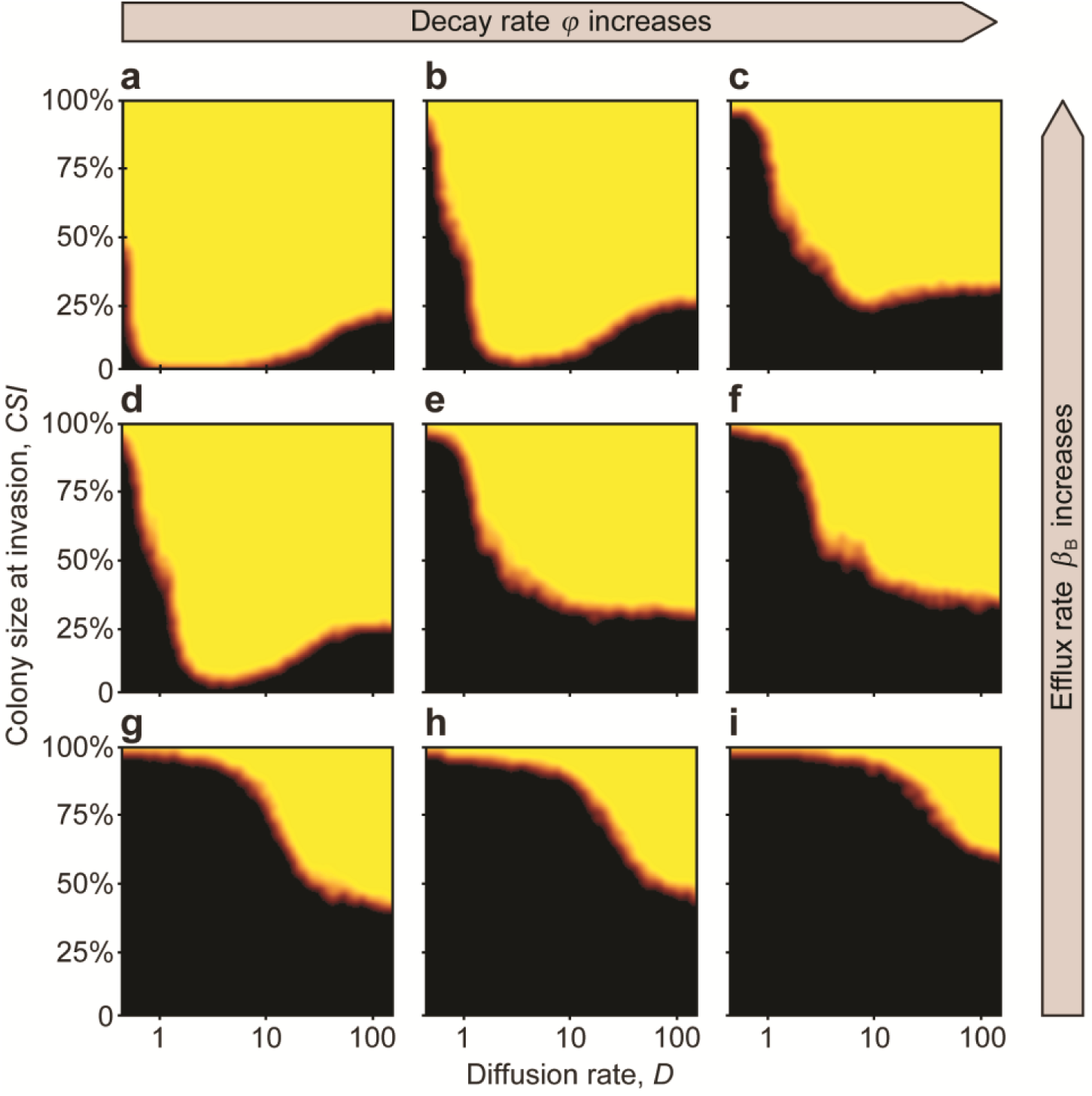
The Minimal Sustainable Colony size (*MSC*) (*Invasion Test 2*). Invasion is initiated when the beneficial-strain colony reaches a defined size (*CSI*) and continues until the habitat is fully colonized by either the beneficial or the parasitic strains. The *MSC* is represented by the orange-red border separating the yellow (**B** wins) and black (**P** wins) regions. From left to right (**a**→**c**, **d**→**f**, and **g**→**i**), the *φ* extracellular decay rate of the antibiotic increases (*φ* = 0.2,0.25,0.3). From top to bottom (**a**→**g**, **b**→**h**, and **c**→**i**), the efflux rate *β*_B_ decreases (*β*_B_ = 1,0.5,0). Simulations were run with 3 replicates for 100 000 generations, or until the population was homogenous. Black areas indicate parameter space where the parasitic strain can invade, yellow indicates parameter space where the antibiotic-producing beneficial strain successfully resists invasion, and orange areas correspond to mixed outcomes. Model parameters are: : *r*_B,0_ = 0.8, *r*_P,0_ = 0.8, *r*_+_ = 0, *c* = 0.4, *ρ*_B,0_ = 1, *ρ*_+_ = 0, *α*_B_ = 0.5, *α*_P_ = 0.5, *β*_P_ = 0, *γ*_B_ = 0.4, *γ*_P_ = 0.4, *D* = 5, *a* = 1, *T* = 1, *k* = 25, *N* = 10 000, *n*_B,0_ = 100, *n*_P,t_ = 10, *f* = 0.01, Δ*t* = 1/10, *u* = 100.

**Figure 4.**
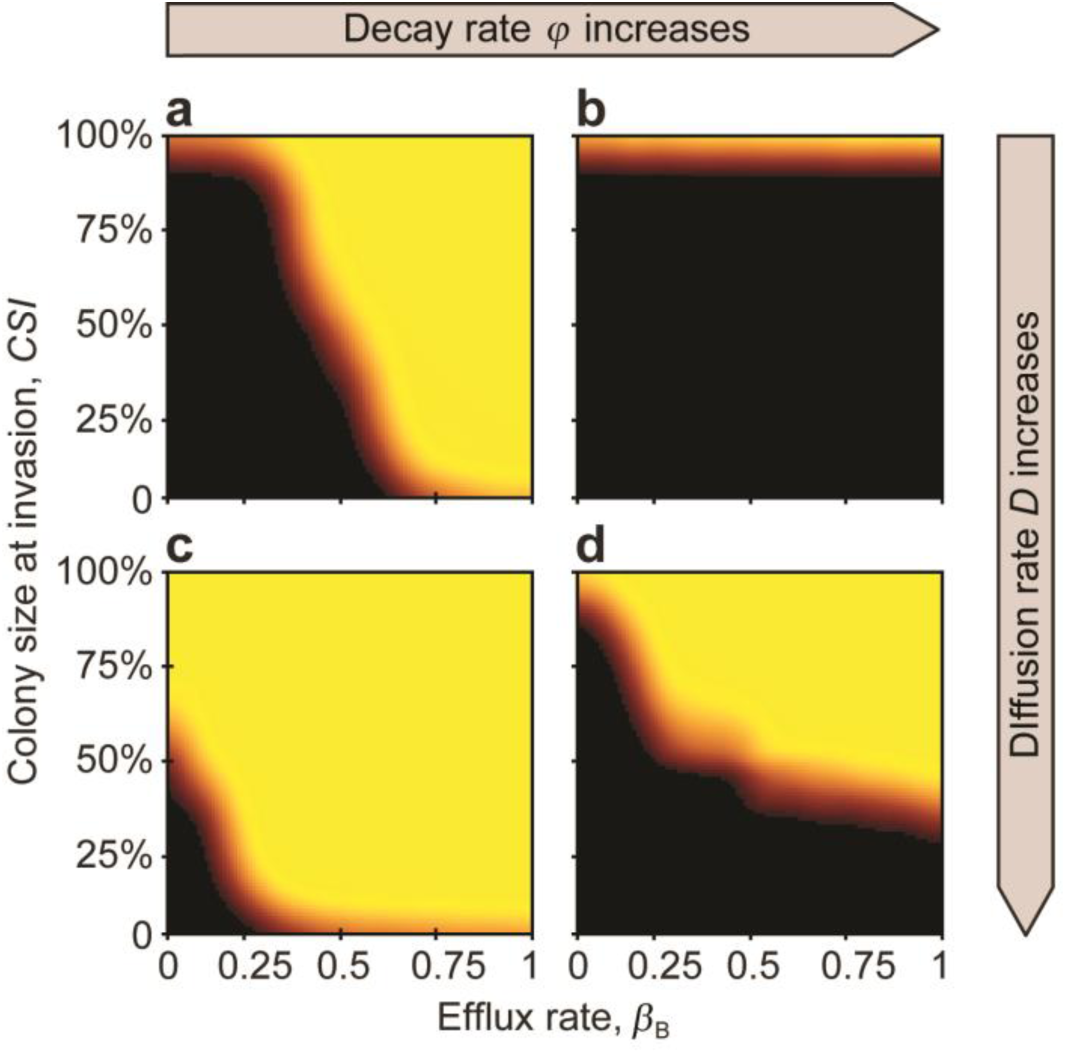
The effect of efflux rate, decay rate, and diffusion rate on the *MSC*. At low diffusion rates (upper row), efflux rate limits the success, while at large diffusion rate (bottom row), colony size is the more limiting factor. From left to right (**a**→**b**, **c**→**d**) extracellular decay rate increases (*φ* = 0.7 and 0.9). From the top to the bottom (**a**→**c**, and **b**→**d**), diffusion rate increases (*D* = 0.5 and 12), respectively. Model parameters are: *r*_B,0_ = 0.8, *r*_P,0_ = 0.8, *r*_+_ = 0, *c* = 0.1, *ρ*_B,0_ = 1, *ρ*_+_ = 0, *α*_B_ = 0.6, *α*_P_ = 0.6, *β*_P_ = 0, *γ*_B_ = 0.3, *γ*_P_ = 0.3, *a* = 1, *T* = 1, *k* = 25, *ρ* = 300, *N* = 10 000, *n*_B,0_ = 100, *n*_P,t_ = 10, *f* = 0.01, Δ*t* = 1/10, *u* = 100.

**Figure 5.**
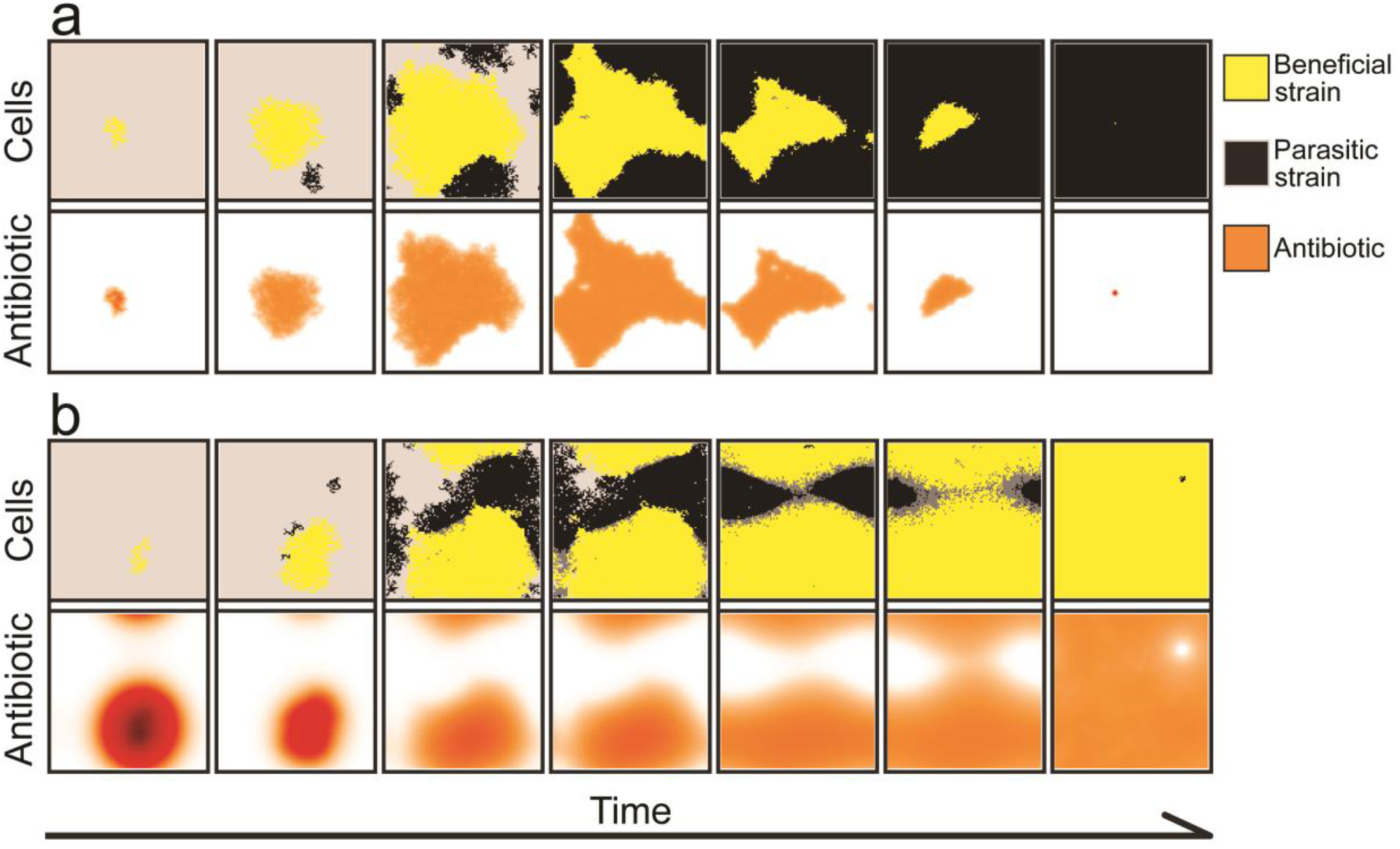
The spatial dynamics for **(a)** low (*D* = 0.5) and **(b)** high diffusion (*D* = 50) rates. **a** A low diffusion rate reduces the protective effect of the antibiotic (orange shading, lower panels), and the parasitic strain (black shading, upper panels) can invade the beneficial strain (yellow shading, upper panels). **b** A high diffusion rate allows the beneficial strain to resist invasion, as considerable amount of antibiotic (lower panels) diffuses beyond the colony boundaries. Antibiotic concentration ranges between zero (white), to intermediate (red-orange), to maximal concentrations (brown-black). Poisoned (cells with *r*_P_ < 0.05) but not yet removed parasitic cells are coloured grey. In these simulations, the beneficial colony was allowed *τ* = 300 time steps to grow before invasion. The snapshots of the simulations are taken every 30 update steps. Model parameters are: *r*_B,0_ = 0.8, *r*_P,0_ = 0.8, *r*_+_ = 0, *c* = 0.4, *ρ*_B,0_ = 1, *ρ*_+_ = 0, *α*_B_ = 0.5, *α*_P_ = 0.5, *β*_B_ = 0.4, *β_P_* = 0, *γ*_B_ = 0.4, *γ*_P_ = 0.4, *φ* = 0.25, *a* = 1, *T* = 1, *k* = 25, *τ* = 300, *N* = 10 000, *n*_B,0_ = 100, *n*_P,t_ = 10, *f* = 0.01, Δ*t* = 1/10, *u* = 100.

The complement to this result is that if the diffusion rate is low, then even a large colony size does not necessarily guarantee success unless the efflux rate is also high enough (Fig. 4a, b). Essentially, if antibiotic efflux is used as the resistance mechanism by the beneficial cells, this can substitute for outright diffusion of the antibiotic, allowing the antibiotic to reach the colony edge, where it can suppress invaders (Fig. 4).

### Non-monotonous effect of diffusion

Interestingly, under some conditions there is a non-monotonous effect of diffusion rate on invasion resistance, such that the Minimum Colony Size (*MSC*) can be much smaller for medium-level diffusion rates. For example, looking at Fig. 3b, for low antibiotic diffusion rates (values 0–1 on the x-axis), the *MSC* is close to 100%; that is, the colony can resist invasion only if more than 95% of the available habitat is already occupied by the producers; otherwise, parasites displace the whole population of antibiotic producers. Similarly, for high diffusion rates (values *D* = 80 – 100 on the x-axis), although smaller, but a considerable colony size still has to be reached. However, the *MSC* curve reaches a minimum between low and high diffusion rates, such that only a 1 – 10% *MSC* is enough to resist invasion (Fig. 3b).

The important result is that for any intermediate efflux and decay parameters and with intermediate diffusion rates, *colonies with practically any non-zero initial size can withstand parasite invasion* (Fig. 3a, b, d). This nonlinearity occurs because, in general, diffusion carries antibiotic to the edge of the antibiotic-producing colony, where it can act against invading **P** strains, but diffusion also carries antibiotic *away from* the edge of the colony. An intermediate diffusion rate turns out to maximise the amount of antibiotic at the fighting front.

## Discussion

The composition of host-associated microbiomes has been shown to correlate with host health status and fitness [4, 81, 82, 83, 84, 85, 86, 87], and thus, there is likely strong selection on host species to evolve mechanisms that favour the assembly of certain kinds of microbiomes over others [11, 12, 27]. Here we have explored how a host can favour the assembly of a defensive microbiome dominated by antibiotic-producing bacteria, hence promoting colonization resistance, in a competitive set-up [7, 23, 74, 88].

We argue that a host can take advantage of an ecological phenomenon known as *bistability*. When two species compete via interference, such as when a bacterial species uses antibiotics to hinder a competitor, the winner of the competition depends partially on the initial population sizes of the two species [9]. If the antibiotic-producer establishes itself with a larger population in the new habitat, it can collectively produce sufficient amount of antibiotic to suppress its competitor and grow until the space of opportunity vanishes for the parasite. In contrast, if the non-producer species starts with the larger or competitively superior population, then the small amount of antibiotic produced by the small colony of the producer is insufficient to suppress the non-producer, hence it wins.

It follows that by using an antibiotic-producer as the initial (or ‘priming’) strain of the microbiome, a host can narrow down the variety of strains able to invade this already established environment [4, 5, 9, 11]. The host is thus efficiently able to canalise the composition of the emerging microbiome. Such priming effects have been demonstrated in various experimental systems [25, 37, 39, 89].

We integrated local interactions and explicit spatial dynamics of cellular and chemical components in our model with the original phenomenological model that laid the foundations if the theory [9]. In this more realistic model, even for large populations, the number of directly interacting cells is relatively modest, and spatial correlations of active agents determine dynamics meaningfully [5, 75]. Furthermore, such an integrated, spatially-explicit model allows us to understand the effect of different antibiotic-resistance mechanisms [73, 76, 77, 78, 79, 80, 90] on the microbiome assembly, and to investigate how attributes of the host surface, which govern the diffusion dynamic of the antibiotic, can modify the outcome. We have also widened the applicability of Scheuring and Yu’s model [9] by reviewing multiple mechanisms allowing a host to prime a defensive microbiome, even if the beneficial can only be recruited from the environment (horizontal transmission), compared to the original model, which made the restrictive assumption that the beneficial strain is strictly vertically transmitted.

We have corroborated the earlier results [9, 13] that antibiotic producers and non-producers can form a bistable system and that the outcome of competition depends on their reproduction rates, how effectively the host is able to selectively promote the beneficial strain, and the initial ratio of the two strains [9]. Once the antibiotic producer is able to gain dominance, in such a system, it can remain dominant for a lifetime, even if the host-provided private resource vanishes or becomes public. The current model also shows that localized interactions do not impede this dominance because the antibiotic itself can diffuse, eventually reaching the colony edge to inhibit invaders. This effect is strengthened when the mode of resistance by the producers is antibiotic efflux.

We also show with the current model that the host resource only needs to remain private for a finite critical time, basically until the beneficial colony reaches a Minimal Sustainable Colony Size (*MSC*), at which point it becomes resistant to a given rate of invasion. The critical time and/or the *MSC* depends on the physiochemical properties of the system, most importantly the decomposition, decay, diffusion, and efflux rates of the antibiotic, and the advantage provided to the beneficial colony by the private resource, all deriving from the fact that colony size determines the amount of antibiotic produced.

Our brief review of the literature suggests that multiple forms of ‘private resources’ exist, including food, space, or host-provided compounds harming undesired strains. Nonetheless, privacy of resources is inherently difficult and costly to achieve, and it is therefore realistic to assume that any host-provided resources will eventually become public. This inevitable transition from private to public, which intuitively might be expected to allow the successful invasion and establishment of parasitic strains, *does not in fact do so*, because of bistability. After a beneficial colony establishes itself, a public resource is in practice only enjoyed by the winner, the beneficial colony.

Finally, we show that an intermediate diffusion rate can maximise the amount of antibiotic accumulating at the colony edge. Our findings suggest that the attributes of the host surface, for example the diffusion rate, can either increase or reduce the effect range of the antibiotic [91]. As there is no conflict of interest between antibiotic-producer and host, their coevolution is expected to optimize the diffusion speed, and hence the effectiveness, of the antibiotic. Overall, evolutionary optimisation can act by minimising the host investment required to attain a beneficial microbiome, by reducing the duration of a private resource supply, and by evolving the optimal physiochemical properties of the habitat, the host surface. If so, then we might also expect that the co-evolution of host and preferred strains results in an efficient and well-conducted build-up of a beneficial microbiome, an orchestrated symbiosis that efficiently narrows down the enormous number of possible scenarios to canalise the emergence of the microbiome towards the most favourable one.

## Acknowledgements

We acknowledge KIFÜ for awarding us access to resource based in Hungary at Budapest, Debrecen, and Szeged. We acknowledge supports from OTKA grants Nr. K100299, and GINOP grant Nr. 2.3.2-15-2016-00057. S.F.W. was funded by a NERC PhD studentship (NERC Doctoral Training Progamme grant NE/L002582/1). D. W. Yu was supported by the National Natural Science Foundation of China (41661144002, 31670536, 31400470, 31500305), the Key Research Program of Frontier Sciences, CAS (QYZDY-SSW-SMC024), the Bureau of International Cooperation project (GJHZ1754), the Ministry of Science and Technology of China (2012FY110800), the University of East Anglia, the State Key Laboratory of Genetic Resources and Evolution at the Kunming Institute of Zoology, and the University of Chinese Academy of Sciences.

## Supplementary information

### Supplementary information 1. The growth dynamic of a colony in the individual-based model

The colony growth follows a logistic growth dynamic in the model (Supplementary Fig. 1). Depending on the choice of *ε*, we observe full colonisation of the surface within a given timeframe. To better investigate the competition dynamics between the two types on a fine timescale, we choose *ε* = 0.01 for our investigations.

**Supplementary Figure 1.**
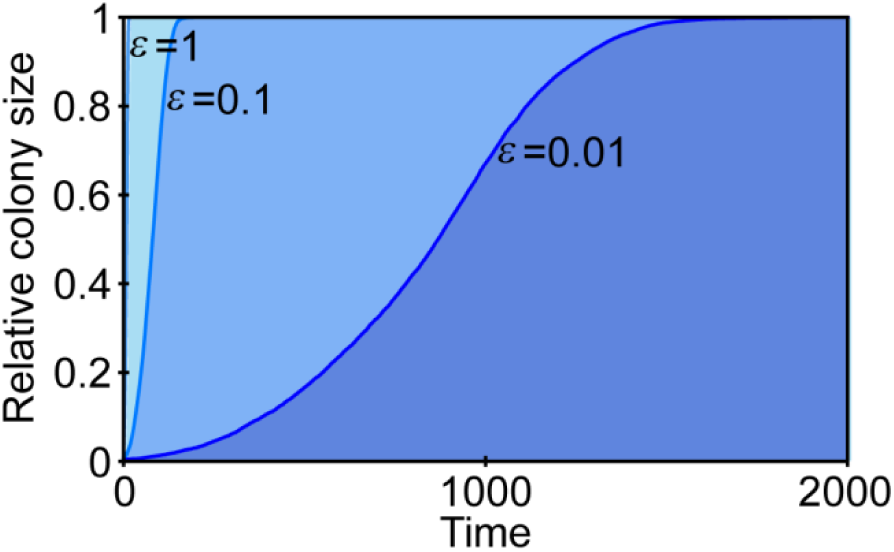
The growth dynamic of a colony follows a logistic trend in the model. We show the relative colony size (y-axis) with respect to time (x-axis) with *ε* = 1 (light blue), *ε* = 0.1 (medium blue), and *ε* = 0.01 (dark blue), where *ε* is the fraction of randomly chosen grid cells that is updated in the cellular reproduction and death processes. The smaller the *ε*, the slower the growth in our model. Relevant model parameters are: D = 5, Δ*t* = 1/10, *u* = 100, for **a** *n*_B,0_ = 100, *N* = 10 000.

### Supplementary information 2. Mathematical formulation of the reaction-diffusion dynamics for the mean-field model

We assume that the antibiotic molecules are point-like particles moving on a host-surface plane. Consequently, we can use reaction-diffusion dynamics to describe change in the extracellular antibiotic concentration *A^Ext^* (**x**,*t*) at points **x** = (*x,y*) (representing the coordinates on a surface) and times *t*

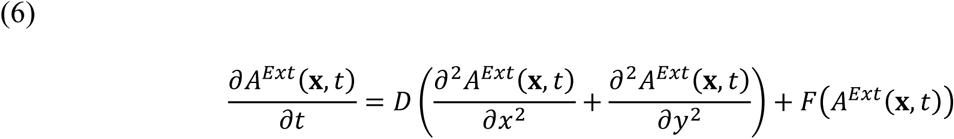

where the first term on the right hand side is the diffusion term, and *F*(*A*^Ext^(**x**,*t*)) is the reaction term, which depends on the extracellular antibiotic concentration (*A^Ext^*) and the positions and types of the cells. Using the above defined parameters and dynamical processes, we can write

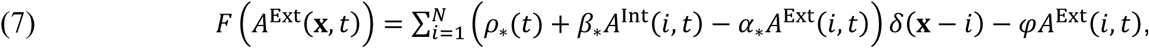

where the antibiotic sources and sinks are summed in the parentheses, *i* is the position of a cell among the *N* cells, which can either be **B** or **P** denoted by * in the bottom index where applicable, *A*^Int^(*i,t*) is the intracellular concentration of the antibiotic at position *i*, and *δ* is the Dirac delta [80].

